# The effect of migration and variation on populations of *Escherichia coli* adapting to complex fluctuating environments

**DOI:** 10.1101/2022.03.22.485294

**Authors:** S Selveshwari, Kaustubh Chandrakant Khaire, Aditee Kadam, Sutirth Dey

**Author notes:** Name and address of the corresponding author*: Sutirth Dey, Biology Division, Indian Institute of Science Education and Research Pune, Dr Homi Bhabha Road, Pune, Maharashtra 411 008, India.

## Abstract

Migration, a critical evolutionary force, can have contrasting effects on adaptation. It can aid as well as impede adaptation. The effects of migration on microbial adaptation have been studied primarily in simple constant environments. Very little is known about the effects of migration on adaptation to complex, fluctuating environments. In our study, we subjected replicate populations of *Escherichia coli*, adapting to complex and unpredictably fluctuating environments to different proportions of clonal ancestral immigrants. Contrary to the results from simple/constant environments, the presence of clonal immigrants reduced all measured proxies of fitness. However, migration from a source population with a greater variance in fitness resulted in no change in fitness w.r.t the no-migration control, except at the highest level of migration. Thus, the presence of variation in the immigrants could counter the adverse effects of migration in complex and unpredictably fluctuating environments. Our study demonstrates that the effects of migration are strongly dependent on the nature of the destination environment and the genetic makeup of immigrants. These results enhance our understanding of the influences of migrating populations, which could help better predict the consequences of migration.

## 1. Introduction

Migration affects several ecological and evolutionary processes, such as a species’ range (Sexton et al., 2009, Kirkpatrick and Barton, 1997, Barton, 2001), composition, and diversity in meta-populations and natural communities (Venail et al., 2008, Albright and Martiny, 2018) and evolution of traits like virulence (Lively, 1999, Boots and Sasaki, 1999) and antibiotic resistance (Perron et al., 2007). Interestingly, when it comes to adaptation, migration can have contrasting effects. For example, migration can impede adaptation in coevolving host-phage systems (Vogwill et al., 2011, Morgan et al., 2005) and microbial communities subjected to warmer temperatures (Lawrence et al., 2016). Similarly, in viruses, migration can reduce the extent of specialization to different tissue types (Cuevas et al., 2003). One of the ways migration negatively affects adaptation is by swamping the destination environment with alleles that are beneficial or neutral at the source environment but maladaptive at the destination (Yeaman and Guillaume, 2009, Kawecki and Holt, 2002, Kawecki and Ebert, 2004). At the same time, several studies have demonstrated that migration can promote adaptation. For example, asexual populations of *Chlamydomonas* exposed to herbicides (Lagator et al., 2014) and yeast populations evolving in the presence of salt stress (Bell and Gonzalez, 2011) adapted more rapidly in the presence of migration. Similarly, Φ6 phage populations adapted better to a novel host when there was migration from populations that had the ability to infect this novel host (Ching et al., 2013). However, when migrants came from control populations unable to infect the novel hosts, there was no effect on the absolute fitness (Ching et al., 2013). Migration can positively influence adaptation by increasing the supply of beneficial mutations, particularly when populations are mutation limited (Holt, 2003, Sexton et al., 2009), such as in asexual microbes.

Interestingly, most empirical studies on how migration influences adaptation in microbes have been carried out only in constant environments, typically in the presence of a single selection pressure (Dennehy et al., 2010, Ching et al., 2013, Lagator et al., 2014, Morgan et al., 2005, Lawrence et al., 2016). However, in nature, organisms are often faced with heterogeneous environments that contain multiple stressors simultaneously. To further complicate matters, the magnitudes of these stresses can fluctuate over time, either predictably or unpredictably. Adaptations in such spatially and/or temporally heterogeneous environments can differ from those in simple constant environments (Levins, 1968, Karve et al., 2016, Cooper and Lenski, 2010, Reboud and Bell, 1997). The effect of migration on adaptation under such complex and fluctuating environments has received less attention (however, see Perron et al., 2007).

Here, we present the results of a study on the effects of different migration rates on adaptation in replicate populations of *Escherichia coli* that were subjected to complex environments undergoing unpredictable fluctuations. We also looked at the effects of migrants that were either clonal or carrying variation. When the immigrants were clonal, the recipient populations evolved reduced fitness compared to the no migration control. Interestingly, the magnitude of fitness reduction varied positively with the fraction of immigrants received. However, treatments that received immigrants with variation in fitness showed little or no change in fitness compared to the no migration control.

## 2. Materials and Methods

For experimental evolution, we used *Escherichia coli* MG1655 with a kanamycin resistance cassette. All cultures were maintained at 37°C and 150 RPM throughout the selection and assays, except where stated otherwise.

### 2.1 Immigrant Populations

This study consisted of two selection experiments. In each selection experiment, we used two populations: the immigrant and the native. The native populations evolved in the complex fluctuating environments and experienced the effects of immigration. The immigrant populations were non-evolving cultures freshly revived every day (see section 2.2 Selection protocol). The two selection experiments differed only in terms of the nature of the immigrant populations. In experiment 1, we used a **<u>C</u>**<u>lonal immigrant population (henceforth C)</u> derived from a single *E.coli* colony and grown in 150ml Nutrient Broth with kanamycin (NB^Kan^) for 18 hours. Multiple 1 ml glycerol stocks (15%) of this culture were prepared and stored at -80°C. In the second experiment, we used the **<u>V</u>**<u>ariation immigrant population (henceforth V)</u>, which was derived from the C population. V population was initiated by reviving 1ml glycerol stock of the C population in 10ml NB^Kan^ followed by inoculation of 1ml of this revived culture in 50ml NB^Kan^. We sub-cultured (1/10th dilution) this population for the next 15 days into 50ml NB^Kan^ every 12 hours. After 15 days (i.e., 30 transfers), 50ml of the grown culture was added to 50ml fresh NB^Kan^ and incubated for another 12 hours. Multiple 1ml glycerol stocks (15%) were prepared and stored at -80°C. The V populations undergo ∼3.3 doublings/transfer which is estimated as log_2_(1 /10^−1^) where 10^−1^ is the bottleneck ratio. Thus, over a period of 15 days (i.e. 30 transfers), the V populations are estimated to have spent ∼100 generations in the benign environment. With an initial population size (N_0_) of 10^8^ cells/ml and final population size (N_f_) of 10^9^ cells/ml and spontaneous mutation rates of the order of 10^−3^ mutation per genome per generation (Lee et al., 2012), we expect ∼3.3×10^6^ mutations to arise in the population, within one transfer. Since this population was maintained with lenient bottlenecks (1/10) at least 10^5^ new arisen mutations are expected to survive the bottlenecking and be carried forward in the subsequent sub-cultures. Furthermore, optimal growth conditions and large culture volumes (N_f_: 10^9^ cells/ml) would also result in clonal interference resulting in a significant number of mutations being maintained in the V population. Thus, we expected the V population to harbour more genetic variation than the C population which were derived from a single colony. To confirm this, we quantified the within-population variance in fitness in the C and V populations using the methodology of an earlier study (McDonald et al., 2012), see supplementary material S1 for further details.

### 2.2 Selection Protocol

We initiated 48 replicate populations from both the C and the V ancestor populations. We then revived 1ml of the corresponding glycerol stocks overnight in 10ml NB^Kan^ and inoculated 20µL of the revived culture (OD_600_ 1.0 – 1.1) into 2 ml of the selection environment

#### Selection environment

The populations were subjected to selection for 30 days in complex environments (i.e., multiple stressors were present simultaneously) and fluctuated unpredictably. We used a selection regimes similar to a previous study (Karve et al., 2015). Briefly, selection involved pH, osmotic (NaCl), and oxidative (H_2_O_2_) stress. We chose the combinations of the stresses so that two of the three components were present at inhibitory concentrations and the third was present at a concentration as found in Nutrient Broth (i.e., pH=7, NaCl=0.5g% and H_2_O_2_=0). We tested several such combinations for their effect on the growth of the WT in a pilot experiment. We chose combinations that resulted in ∼ 40 – 70% reduction in growth compared to NB growth. We chose a total of 28 combinations and a random sequence of 30 environments from a uniform distribution with replacement (see supplementary Table S2 for full list). At the end of 30 rounds of selection, populations were estimated to have undergone selection for a maximum of ∼200 generations (∼6.67 doublings/transfer (i.e., log_2_(100)) × 30 days). While the growth rates of populations in the different sub-lethal environments were not the same, all treatments were exposed to the same sequence of environments. This ensured that all populations experienced similar generation times over the 30 rounds of selection.

#### Migration treatment

We used four levels of migration, namely 0% (control), 10% (low migration), 50% (intermediate migration) and 90% (high migration), with 12 replicate populations per migration treatment. If we simply added the immigrants to the native population, the total population size would increase with the migration rate, thus resulting in large differences in population sizes across treatments. Previous studies have shown that population size can affect evolutionary outcomes in microbial populations (Chavhan et al., 2019). Therefore, to avoid the confounding effect of population size, we kept the inoculum size constant (∼10^7^ cells) and defined migration as the percentage of immigrants in the inoculum. For example, in the low (10%) migration treatment, 10% of the individuals in the subculture inoculum consisted of immigrants. The remaining 90% were individuals from the native population, evolving under a complex fluctuating environment. The proportions were adjusted based on the OD_600_ values of the immigrant and native populations at the time of subculture. When OD_600_ = 1, the culture contained ∼10^9^ cells/ml of NB (S Selveshwari, personal observations).

Every day, we revived 1ml glycerol stock of the C or V population in 10ml NBKan. OD_600_ of the revived culture was adjusted to 1.0 – 1.1 and used as immigrants. We also measured OD_600_ of the native populations and used an appropriate culture volume containing the required inoculum size for the sub-culture. When the OD_600_ of these populations was less than 0.3, the populations were considered extinct, and the native population was obtained from the previous non-extinct population, stored at 4°C. The threshold of 0.3 was used since populations with growth below this OD failed to survive subsequent rounds of selection (S Selveshwari, personal observation). We stored the selected populations as glycerol stocks at the end of 30 days.

### 2.3 Fitness assays

We measured the fitnesses of the evolving populations during and after selection. We noted the OD_600_ in the selection environment every 24 hours (i.e., at carrying capacity, just before sub-culturing). We used the geometric mean of these values over the 30 days of selection to measure fitness under fluctuating stress (Orr, 2009). Using the same data, we also estimated the probability of extinction (OD_600_ < 0.3) during selection.

We measured post-selection fitnesses as growth rate and yield of the populations in three representative environments: environment 1: pH 8.5+salt 4.5g%; environment 2: pH 5+0.5µL H_2_O_2_; environment 3: salt 2.5g%+0.7µL H_2_O_2_. All 48 replicate populations of each selection experiment were assayed twice in each assay environment. 4µL of glycerol stock was revived in 2ml NB^Kan^, overnight. We measured the OD_600_ of the revived culture and inoculated a volume containing 10^7^ cells (assuming 10^9^ cells/2ml when OD_600_ = 1) in 2ml assay environments in 24-well tissue culture plates. The OD_600_ was measured every 2 hours for 24 hours at 37°C and slow continuous shaking using a plate reader (Synergy HT BioTek, Winooski, VT, USA). We used population growth rate and yield as fitness measures (Chavhan et al., 2019, Karve et al., 2015). The growth curve data were analyzed using a custom python script that fits overlapping straight lines over 6-hour windows. We estimated growth rate as the maximum slope of the curve and yield as the maximum OD_600_ reached in 24 hours.

### 2.4 Statistical analysis

All fitness comparisons were performed independently for the two selections. The geometric mean of growth during selection was compared across the migration treatments (each with 12 replicate populations) using separate one-way ANOVAs with migration treatment (0, 10, 50, 90) as a fixed factor. Fitness measured as extinction probability was analyzed similarly however, since the data was in fractional form, they were subjected to arcsine-square root transformation prior to ANOVA (Zar, 1999). We compared post-selection fitnesses (growth rate and yield) for each assay environment, using separate two-way mixed model ANOVAs. We treated migration treatment (0, 10, 50, 90) as a fixed factor. The growth rate and yield from the two rounds of assays were considered measurement replicates. The biological replicates (12 levels) acted as random factors and were nested in migration treatment. To account for the inflation of family-wise error rate, the *P*-values of the main effect of migration were subjected to Holm–Šidák correction (Abdi, 2010). We performed pairwise comparisons using Tukey’s post hoc analysis when the corrected *P*-values was significant. We also computed the Cohen’s *d* statistics (Cohen, 2013) to measure the effect sizes (Sullivan and Feinn, 2012). The biological significance of the differences between the treatments were interpreted as small, medium and large for 0.2 < *d* < 0.5, 0.5 < *d* < 0.8 and *d* > 0.8, respectively.

We performed all the ANOVAs on STATISTICA v7.0 (Statsoft Inc.). Cohen’s *d* statistics were estimated using the freeware Effect Size Generator v2.3.0 (Devilly, 2004).

## 3. Results

### 3.1. Clonal immigration impedes adaptation in complex and unpredictable environment

After 30 rounds of clonal migration and selection (corresponding to ∼200 generations in the no migration control populations), we found that the control populations (i.e., those that did not receive any immigrants) adapted significantly more than populations that received immigrants (from clonal source, C). The geometric mean of growth during selection was significantly different between the migration treatments (Fig. 1A; *F*_3,44_ = 28.76, *P* = 1.88E-10). The no migration control populations had the highest GM of growth. However, this was only significantly higher than intermediate (Tukey’s p = 0.01; Cohen’s *d* = 1.49 (large)) and high (Tukey’s p = 0.0002; Cohen’s *d* = 3.56 (large)) migration treatments. The reduced GM of growth of high migration treatment was also significant w.r.t low (Tukey’s p = 0.0002; Cohen’s *d* = 2.88 (large)) and intermediate (Tukey’s p = 0.0002; Cohen’s *d* = 2.42 (large)) migration treatments. This reduction in growth was accompanied by a significant effect in terms of extinction probability (Fig. 1B; *F*_3,44_ = 13.96, *P* = 1.56E-06). Tukey’s post hoc indicated that the high migration treatment had significantly greater extinction probability compared to all treatment population (control: p = 0.0002; Cohen’s *d* = 2.10 (large); low: p = 0.0002; Cohen’s *d* = 2.09 (large); intermediate: p = 0.004; Cohen’s *d* = 1.72 (large)). Thus, during selection, the presence of migration reduced the populations’ ability to survive and grow in the selection environment, resulting in a significant increase in extinction probability at high migration. We next tested how this reduced survivability and growth affected the overall adaptation.

**Figure 1.**
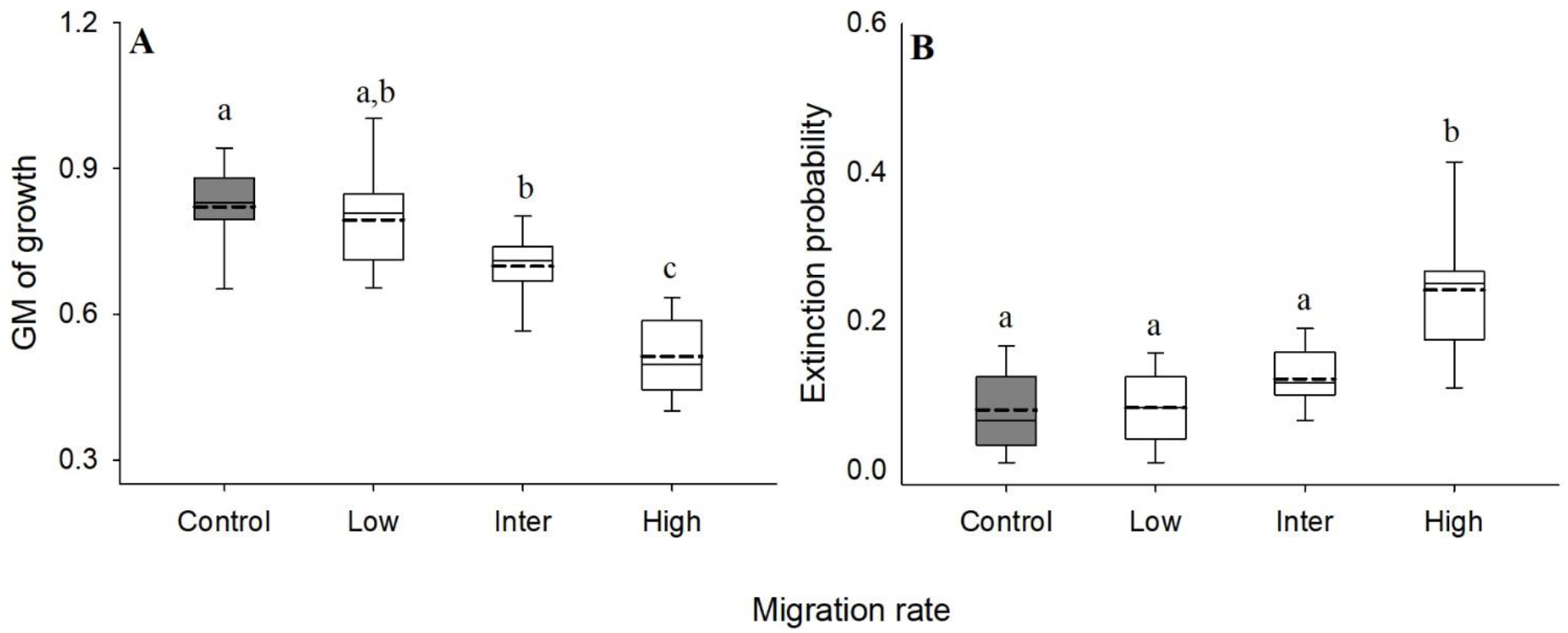
Effect of clonal migration during selection in complex, unpredictable environments. Fitness was measured as A) Geometric mean of growth during selection. B) Probability of extinction in the selection environments, during selection. Each box plot represents data from 12 replicate populations. Solid lines represent median, dotted lines denote mean, whiskers denote 10^th^ and 90^th^ percentiles, and dots denote 5^th^ and 95^th^ percentile. Box plots denoted by different letters are significantly different from each other (P < 0.05 in Tukey’s posthoc analysis).

### 3.2. Populations receiving clonal immigrants evolved lower fitness than control populations

After selection, we estimated the extent of adaptation as growth rate and yield in three complex environments. When the fitnesses across all three environments were analyzed together in a single ANOVA, we saw that the interaction between migration treatments and assay environment was significant (Fig. 2; Growth rate: *F*_6,144_ = 2.43, *P* = 0.03; Yield: *F*_6,144_ = 4.62, *P* = 4.0E-04). A significant interaction effect denoted that the fitness difference between the migration treatments varied depending on the choice of the complex environment.

**Figure 2.**
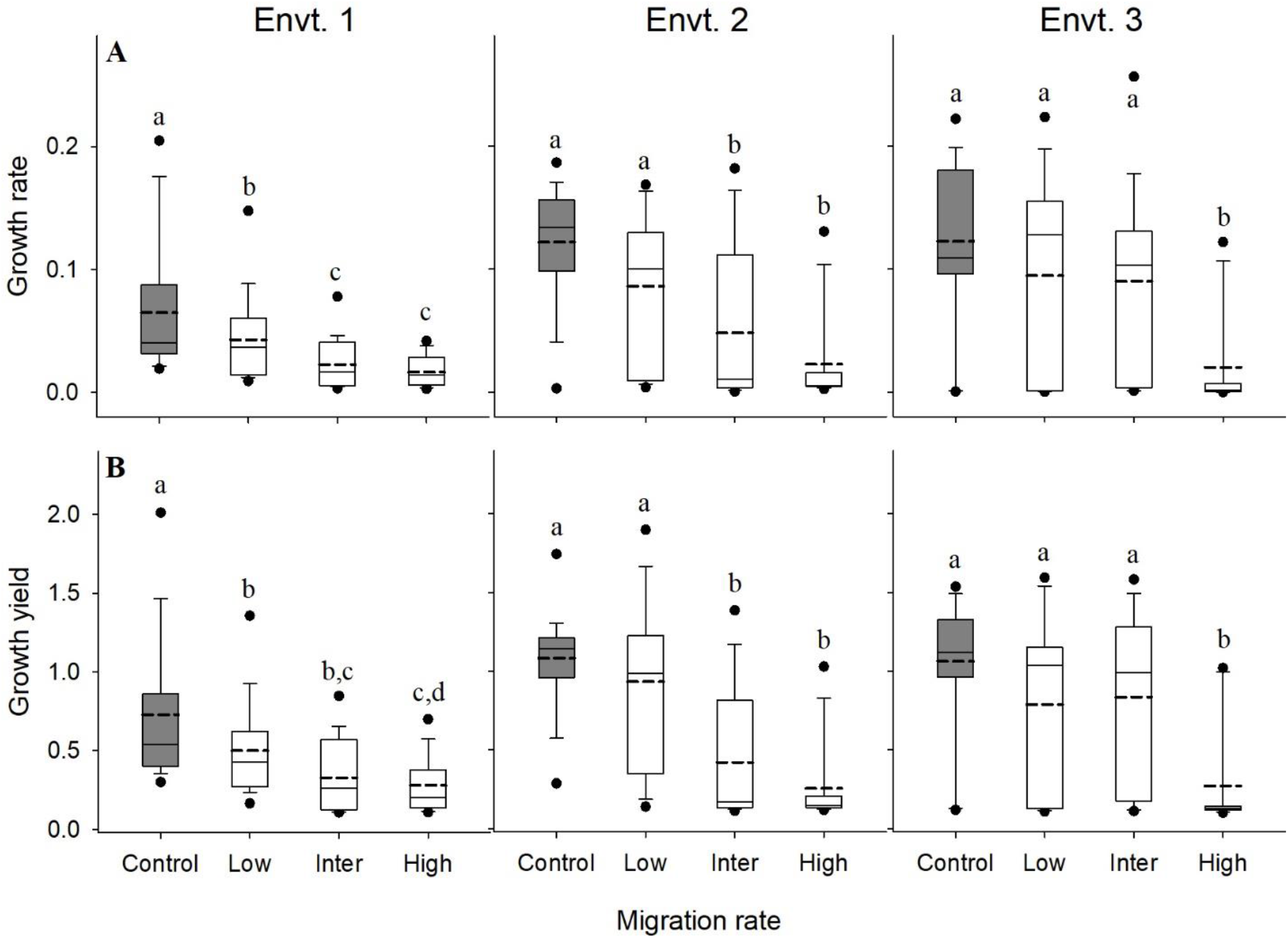
Effect of clonal migration on post-selection fitness. Two fitness proxies, A) growth rate and B) growth yield, were measured in three complex environments. See methods for the composition of the three complex environments. Each box plot represents 24 values, i.e., 12 replicate populations, assayed twice. Solid lines represent median, dotted lines denote mean, whiskers denote 10^th^ and 90^th^ percentiles, and dots denote 5^th^ and 95^th^ percentile. Box plots denoted by different letters are significantly different from each other (P < 0.05 in Tukey’s posthoc analysis).

Therefore, the effect of clonal migration on fitness in the three complex environments was analysed independently. The results of ANOVAs, *P*-values and Cohen’s *d* of all pairwise comparisons are summarized in supplementary tables S3 and S4. Briefly, all migration treatments had significantly lower growth rates and yield than the no migration control populations in environment 1 (pH 8.5+salt 4.5g%). The growth rate of the low migration treatment was significantly higher than both intermediate and high migration treatments, which were not significantly different from each other. Similarly, although the yield of the low migration treatment was higher, it was statistically significant w.r.t only the high migration treatment. However, both intermediate and high migration treatments were not significantly different. In environment 2 (pH 5+0.5µL H_2_O_2_), only two migration treatments (intermediate and high) had significantly lower growth rates and yield w.r.t the control. However, the growth rate of the low migration populations was only marginally insignificant (Tukey’s p = 0.054) w.r.t control populations in environment 2. Both the growth rate and the yield of the low migration treatment were significantly higher than the intermediate and high migration treatments. However, there was no significant difference between growth rates and yields of intermediate and high migration treatments. In environment 3 (salt 2.5g%+0.7µL H_2_O_2_), the growth rate and the yield of only the high migration treatment were significantly lower than all other treatments. Thus, taken together, we saw that migration can have an overall negative effect on evolutionary outcomes under complex and unpredictable environments.

### 3.3. Immigrants carrying variation has little effect of fitness during selection

When the migrant populations carried variation (V population) (see suppl. material S3 for evidence of greater variation in V population), migration had an effect only at high levels of migration. A significant main effect was observed w.r.t geometric mean of growth (Fig 3A; *F*_3,44_ = 34.14, *P* = 1.48E-11). GM of growth was reduced at high levels of migration and this reduction was significantly different from all other treatments in Tukey’s post-hoc analysis (control: p = 1.69E-04, Cohen’s *d* = 2.25 (large); low: p = 1.69E-04, Cohen’s *d* = 3.37 (large); intermediate: p = 1.69E-04, Cohen’s *d* = 3.52 (large)). Subsequently, extinction probability also had a significant main effect of migration (Fig 3B; F_3,44_ = 6.07, p= 0.0015) and high migration treatment had an elevated probability of extinction. However, this increase in extinction was significantly different from only low and intermediate migration treatment (low: p = 9.32E-04, Cohen’s *d* = 2.02 (large); intermediate: p = 0.03, Cohen’s *d* = 0.90 (large)). Thus in contrast to migration from a clonal source, the negative effect of migration, during selection, was diminished when the migrants carry variation.

**Figure 3.**
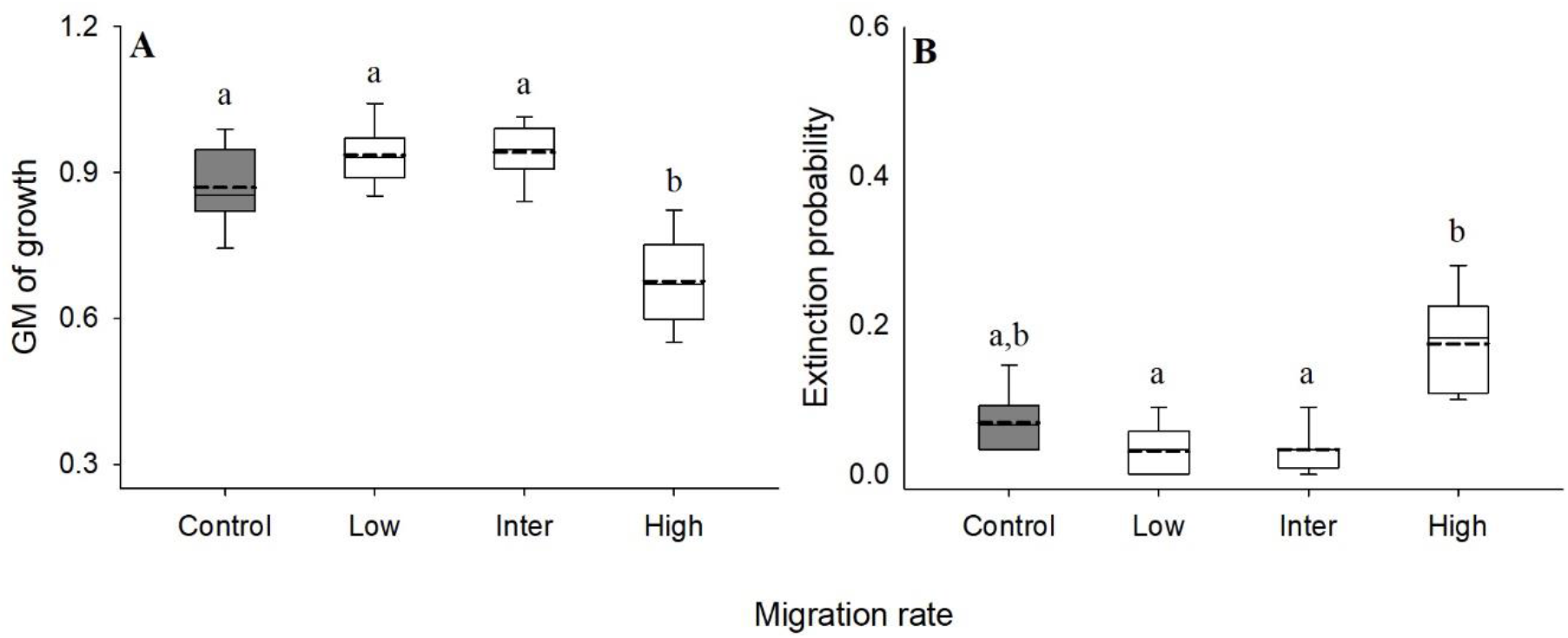
The effect of migration, when the immigrants carry variation. Fitness was measured as A) Geometric mean of growth during selection. B) Probability of extinction in the selection environments, during selection. Each box plot represents data from 12 replicate populations. Solid lines represent median, dotted lines denote mean, whiskers denote 10^th^ and 90^th^ percentiles, and dots denote 5^th^ and 95^th^ percentile. Box plots denoted by different letters are significantly different from each other (P < 0.05 in Tukey’s posthoc analysis).

### 3.4. Presence of variation in the immigrant pool counters the negative effect of migration

Following section 3.2, we analyzed the fitnesses of the populations receiving migrants with variation in the three complex environments using independent ANOVAs (see suppl. Tables S5 and S6 for summary of *P* and Cohen’s *d* values). In contrast to clonal migration, we found that the main effect of migration was either non-significant (growth rate in environments 2 (pH 5+0.5µL H_2_O_2_) and environment 3 (salt 2.5g%+0.7µL H_2_O_2_)) or, when significant, the effect was limited to only the populations receiving a high level of migration. The growth rate and yield in environment 1 (pH 8.5+salt 4.5g%) were significant only between the no-migration control and high migration treatments. The main effect of growth rate in environment 2 was not significant (*F*_3,44_ = 1.9, *P* = 0.144). However, yield in environment 2 was significant, and Tukey’s post hoc analysis showed that high migration treatment had significantly lower yield than control and low migration treatments (control: p = 0.009, Cohen’s *d* = 0.934 (large); low: p = 0.007, Cohen’s *d* = 0.922 (large)). Like environment 2, growth rate in environment 3 was also not significant (*F*_3,44_ = 2.56, *P* = 0.13). Similarly, yield in environment 3 was significant, and Tukey’s post hoc analysis showed that high migration treatment has a significantly lower yield than only no-migration control (control: p = 0.003, Cohen’s *d* = 1.057 (large)). These results illustrate that variation in the migrant pool can ameliorate the negative effects of migration.

## 4. Discussion

In this study, we investigated the effect of immigration on the adaptation of asexual populations in complex and unpredictably fluctuating environments. Immigration from a clonal and non-evolving source (ancestor) population resulted in a reduction in fitness during selection (Fig. 1) as well as post-selection (Fig. 2). During selection, as the proportion of immigrants increased, the geometric mean (GM) of growth decreased (Fig. 1A), and the extinction probability increased (Fig: 1B). After ∼200 generations, the populations that received immigrants during selection had adapted less with a reduced growth rate (Fig. 2A) and yield (Fig. 2B). The fitness reduction increased with the fraction of immigrants in the evolving populations.

These results are in apparent contradiction with several studies where migration promotes larger and/or faster adaptation in asexual organisms (Bell and Gonzalez, 2011, Lagator et al., 2014). In particular, it disagrees with a previous study (Perron et al., 2007) that used a clonal source population as source of migration and showed that increasing immigration rates leads to rapid evolution. In their study, although the rates of adaptation were faster in benign (single antibiotic) environments than in harsh (two antibiotics) environments, the effect of migration was positive in all cases. In contrast, we saw increasing adverse effects of migration with increasing migration rates. One possible reason for such contrasting effects could be attributed to the choice of migration rates used in Perron *et al*. (Perron et al., 2007) and our study. However, we note a key difference in the definition of migration rate between the two studies. Perron *et al*. (Perron et al., 2007) have defined migration rate as proportion of stationary phase culture resulting in migration rates that are 0%, 0.1%, 1%, and 10% of ancestral culture. However, if these numbers were translated to proportion of immigrants in the inoculum (according to the definition used in our study), the migration rates used in Perron *et al*. (Perron et al., 2007) are similar to our treatment; i.e., 0%, 10%, 50%, and 90% (see supplementary material S2 for details). The other possible reason for the discrepancy in results, despite similarities in source population and migration treatments, could be the stress intensity used in the selection environments. While both experiments had multiple stress components, Perron *et al*. (Perron et al., 2007) used lethal concentrations of stress, whereas we used sub-lethal concentrations. Lethal stresses create a sink environment where population size is expected to decline without sustained migration (as pointed out by the authors themselves) (Holt and Gomulkiewicz, 1997, Holt, 1997, Dias, 1996). As in the case of the two experiments (Perron et al., 2007) and this study), immigration from a clonal source population can be expected to promote adaptation in terms of its demographic effect, i.e., changes in population sizes, which in turn can influence the adaptive dynamics of these populations. However, in non-lethal/ non-sink environments, like in our experiment, migration provided no demographic advantage as populations can persist here without immigration. Instead, the effect was largely negative as selection for locally fitter individuals can be diluted by increasing proportions of immigrants (Kawecki and Ebert, 2004, Kawecki and Holt, 2002, Lenormand, 2002). The two studies taken together highlight that the nature of the environment faced by the evolving populations needs to be considered when studying the effect of migration on adaptation. The results can be very different when populations evolve in sub-optimal environments compared to those evolving in lethal environments.

Since clonal immigrants in sub-optimal environments did not provide any significant advantage to adaptation, we next investigated how variation in the immigrants influences adaptation to complex and unpredictable environments. To this end, we conducted a second selection experiment using a source population with a larger variation in fitness (Fig. 3). Immigration from this variant population had little or no effect on the evolving population, as seen from fitness measured during and post-selection (Figs. 4 and 5). Populations receiving low and intermediate levels of migration did not show a reduction w.r.t any aspects of fitness. However, a reduction in fitness was observed when the populations were subjected to high migration.

**Figure 4.**
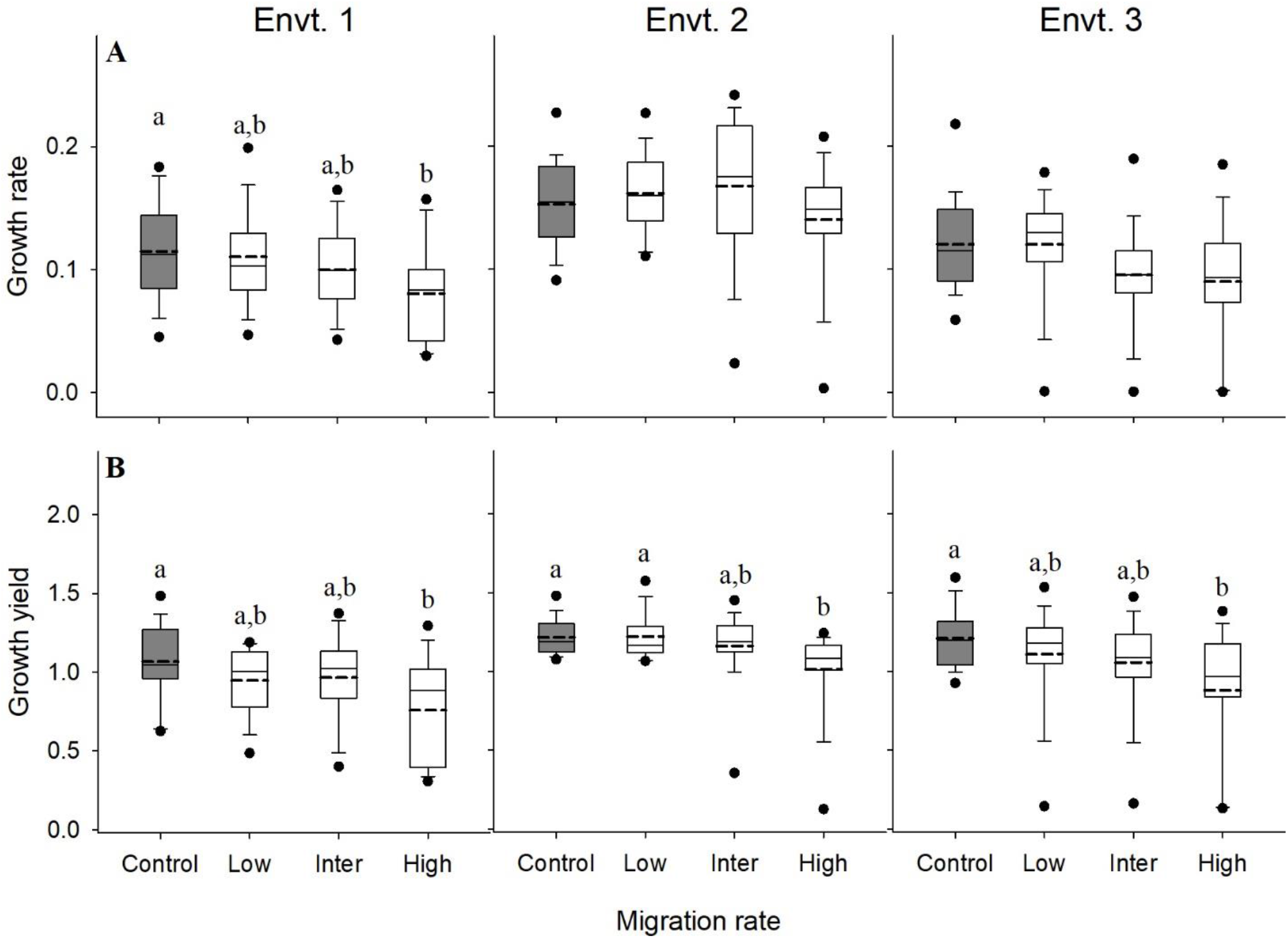
Effect of migrants with variation on post-selection fitness. Fitness was estimated as A) growth rate and B) yield in three complex environments. Each box plot represents data of 24 values, i.e., 12 replicate populations, assayed twice. Solid lines represent median, dotted lines denote mean, whiskers denote 10^th^ and 90^th^ percentiles, and dots denote 5^th^ and 95^th^ percentile. Box plots denoted by different letters are significantly different from each other (P < 0.05 in Tukey’s posthoc analysis).

It has been previously shown in bacteriophages that increased variation due to migration can promote adaptation (Dennehy et al., 2010). However, in that study, the positive effect was limited to immigration from source populations grown in the same environment as the selection environment. Migration between similar environments could have promoted the spread of beneficial variation between sub-populations (Kassen, 2014). On the other hand, in our study, the source population was grown in a benign environment, unrelated to the selection environment. Migration from such a source is not limited to only beneficial variants.

Additionally, the distribution of fitness effects of new mutations is expected to vary greatly in a complex and fluctuating environment with rugged and shifting fitness landscapes (Van Cleve and Weissman, 2015). Thus, in non-sink but complex unpredictable environments, we saw that the benefit of increased variation was only enough to counter the adverse effects of migration. These results agree with theoretical predictions that variation in the migrant pool can ameliorate the harmful effects of migration (Barton, 2001). However, if the variation is too high, one can expect adverse effects on fitness, including extinctions (Barton, 2001). Significant reduction in all aspects of fitness with high migration can indicate the existence of such a limit on the positive effects of increased variation via migration.

Populations receiving high levels of migrants, from both clonal and variant sources, relied on recurrent immigration and revival from previous time points for survival under complex, unpredictable conditions. This was a clear demonstration of the creation of pseudo-sinks where viable environments appear to have become sink environments due to high migration (Watkinson and Sutherland, 1995). Repeated introduction of individuals into sub-optimal environments can limit adaptation and result in the populations being in a constant state of maladaptation, a phenomenon commonly observed at range margins (Kirkpatrick and Barton, 1997).

## 5. Conclusions

Migrating individuals play multiple roles (demographic, variation) in the destination environment (Garant et al., 2007). The relative importance of these aspects of migration and their influence on adaptation is dependent on the quality of the environment. Maintaining a sustainable population might be more critical in a lethal sink environment. However, the supply of variation is more critical in sub-optimal environments, without which migration can create pseudo-sink environments. Since sub-optimal environments are likely more prevalent in nature, studies conducted under such conditions need to understand the effects of migration fully. Additionally, it becomes vital to consider the interactions between the role of migrating individuals and the environment into which they migrate.

## Acknowledgements

We thank Gayathri Pananghat and Milind Watve for their valuable inputs. SS was supported by a Junior Research Fellowship initially sponsored by IISER Pune and then by Department of Biotechnology (DBT), Govt. of India. AK was supported through INSPIRE fellowship of Department of Science and Technology (DST), Government of India. This project was supported by a grant from Department of Biotechnology, Government of India (#BT/PR22328/BRB/10/1569/2016) and internal funding from IISER Pune. The authors declare no conflict of interest.

## Supplementary material

### Supplementary material S1. Quantification of within-population variance between C and V populations

To quantify the variation accumulated in the V population, we assayed the fitnesses of 72 individual colonies of both C and V populations in 6 environments. We used the within-population fitness variance as a proxy for genetic variation (McDonald, Hsieh et al. 2012). 1ml glycerol stock of the C or V population was revived in 10ml NB^Kan^, overnight. The revived cultures were streaked on nutrient agar (NA) with kanamycin from where, we made single colony suspensions from 6 similar-sized colonies of both C and V populations. We then inoculated 10µL of the single colony suspension in 200µL of six assay environments, assayed in a single 96-well plate. The environments included sub-lethal concentrations of all stress combinations and NB^Kan^. Environment I: pH 5+Salt 3.5g%, Environment II: pH 8.5+Salt 2.5g%, Environment III: pH 5+4µl 0.3% H_2_O_2_, Environment IV: pH 8.5+2µl 0.3% H_2_O_2_, Environment V: Salt 2.5g%+1.5µl 0.3% H_2_O_2_ and Environment VI: NB^Kan^. H_2_O_2_ was added 2 hours after inoculation, where required. The populations were continuously monitored (OD_600_) for 24 hours, from inoculation, using a plate reader (Synergy HT) at 37°C and continuous medium shaking. We repeated the entire procedure over 12 days for a total of 72 colonies from each of the C and V populations. Growth rate and yield were estimated using a custom python script; see section 2.3 for details. Within-population variance in fitness was measured as the coefficient of variation (CV) in fitness of the C and V populations. The CV was computed between the six colonies assayed every day. CV was computed for fitness measured as both growth rate and yield. The CV of the C and V populations were compared in each of the six environments using paired t-Test. The inflation in family-wise error rate was controlled using Holm–Šidák correction.

**Figure.**
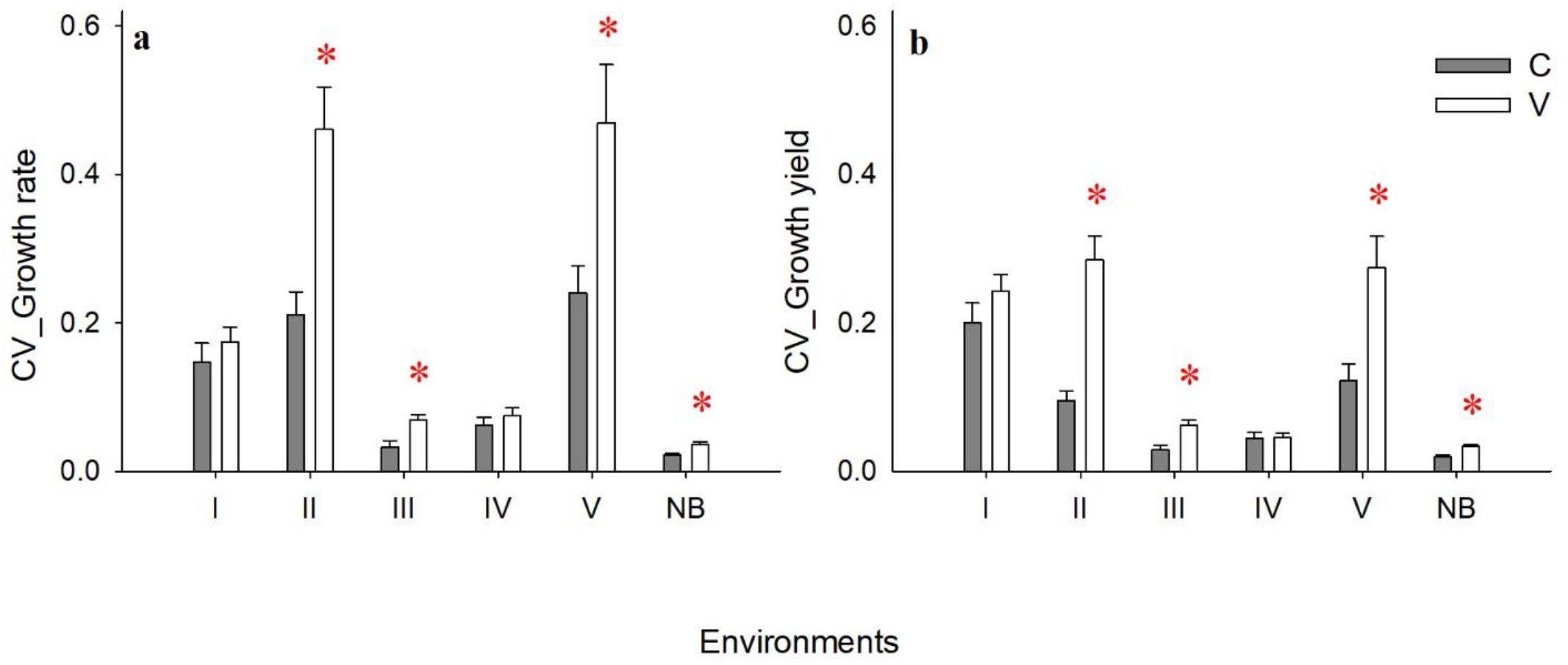
The average coefficient of variation of the clonal and variant populations. We measured the coefficient of variation (CV) of growth rate and yield of the two populations, assayed in 6 representative environments. Each bar represents the average CV over 72 individual colonies assayed over 12 days. Error bars are SE of the mean. * denote *P*-value < 0.05 in paired t-Test after Holm-Šidák correction.

After ∼100 generations of lenient (1/10) bottlenecks, in a benign environment, the V population showed a significant increase in variation in fitness, measured as coefficient of variation, in 4 out of 6 tested environments (Fig. S1 and Table S2). The increase in variation was seen w.r.t both growth rate and yield in environments II, III, V and NB. Thus, the V population had a larger within-population fitness variance than the C population and was used as the source of migrants in a second selection experiment.

## Supplementary material S2. Defining migration rates

In the literature on migration in bacteria, there is some difference in terms of how migration rates are computed. For example, Lagator *et al*. (2014) define migration rate as the proportion of immigrant cells in the inoculum (cells transferred into fresh media) whereas, Perron *et al*. (2007) define migration rate as “the proportion of bacterial cells transferred from a fresh stationary phase culture of the ancestral clone grown overnight in unsupplemented KB to the selection”.

We show below that if one uses a consistent definition, then the rates we investigated are close to what the earlier studies have used, which would allow for qualitative comparisons of the evolutionary outcomes under different scenarios. In our study, we have defined migration rate as the proportion of immigrants in the inoculum which is similar to Lagator *et al*. 2014 who report that their lowmig and highmig treaments contained 55% and 80% cells from the source at first transfer, respectively. When we translate the migration rates from Perron *et al*. (2007) into proportion of migrants in the inoculum, their migration rate are comparable with our study. See the following table for detailed calculation using the data given in Perron *et al*. (2007).

| A: Treatment from Perron et.al., 2007 | B: Resident cells | C: Migrants cells | D: Total inoculum size (B+C) | E: Proportion of migrants in inoculum (C/D) |
| --- | --- | --- | --- | --- |
| 0% | 1.00E+06 | 0.00E+00 | 1.00E+06 | 0.00% |
| 0.10% | 1.00E+06 | 1.00E+05 | 1.10E+06 | 9.09% |
| 1% | 1.00E+06 | 1.00E+06 | 2.00E+06 | 50.00% |
| 10% | 1.00E+06 | 1.00E+07 | 1.10E+07 | 90.91% |

As can be seen from column E of this table, the percentages used by Perron *et al*. (0, 9.09, 50.0 and 90.91) are actually very close to what we have used (0, 10, 50 and 90 respectively).

---

**Table S1.** Summary of the *P*-values of paired T-test of coefficient of variation in fitness between C and V ancestors, after Holm-Šidák correction.

|  | *P*-value after Holm-Šidák correction |  |
| --- | --- | --- |
| Environment | Growth rate | Yield |
| I | 0.607 | 0.371 |
| II | 0.008 | 0.0004 |
| III | 0.005 | 0.0004 |
| IV | 0.471 | 0.852 |
| V | 0.017 | 0.005 |
| NB | 0.048 | 0.012 |

**Table S2.** Sequence of complex fluctuating environments used during selection.

| **Complex fluctuating environment** |  |  |  |
| --- | --- | --- | --- |
| **Day** | **pH** | **NaCl (g/100ml)** | **H<sub>2</sub>O<sub>2</sub> (μL)** |
| 1 | 5 | 4 | - |
| 2 | 4.5 | 3 | - |
| 3 | 5 | 4.5 | - |
| 4 | - | 2.5 | 0.8 |
| 5 | 5 | 3.5 | - |
| 6 | 5 | - | 0.6 |
| 7 | 5 | 3 | - |
| 8 | - | 4 | 0.5 |
| 9 | - | 3 | 0.5 |
| 10 | 5 | 3 | - |
| 11 | - | 4 | 0.5 |
| 12 | - | 3 | 0.5 |
| 13 | 5 | - | 0.7 |
| 14 | - | 3 | 0.5 |
| 15 | 8.5 | - | 0.5 |
| 16 | - | 3 | 0.7 |
| 17 | 5 | - | 0.5 |
| 18 | 5 | 3 | - |
| 19 | 9 | - | 0.5 |
| 20 | 5 | 5 | - |
| 21 | - | 3 | 0.5 |
| 22 | 9 | - | 0.5 |
| 23 | 8.5 | 5 | - |
| 24 | 4.5 | 3 | - |
| 25 | 8.5 | 4 | - |
| 26 | - | 2 | 0.8 |
| 27 | 5 | 3 | - |
| 28 | 8.5 | 4 | - |
| 29 | 5 | 4 | - |
| 30 | 8.5 | 4 | - |

**Table S3.**
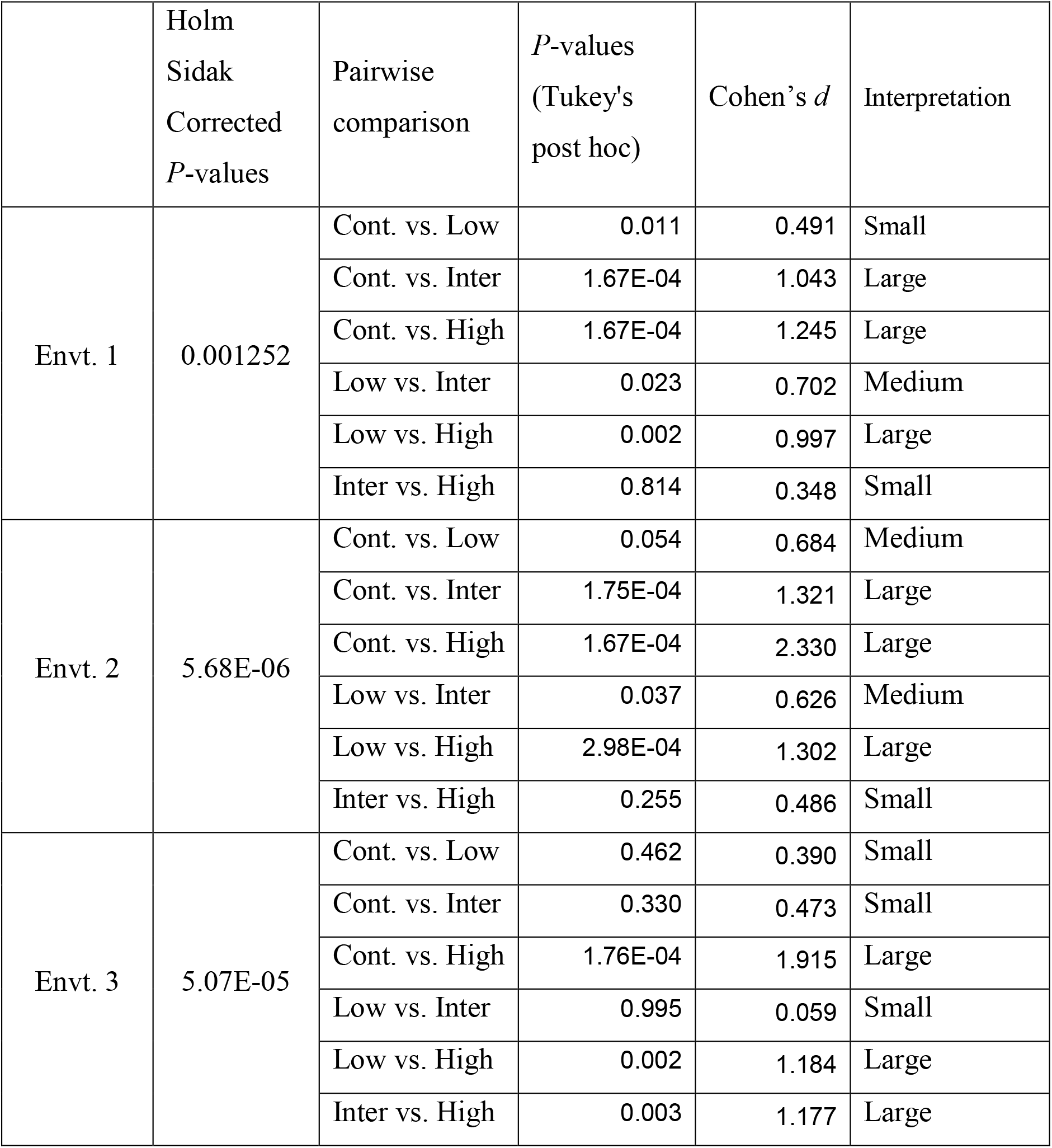
Summary of the *P*-values for the growth rate of clonal migration post-selection. *P*-values of the main effect of migration was reported after Holm-Šidák correction for familywise error. Migration treatments were compared with each other using Tukey’s post-hoc analysis. Pairwise effect sizes were computed as Cohen’s *d*.

|  | Holm<br>Sidak<br>Corrected<br>*P*-values | Pairwise<br>comparison | *P*-values<br>(Tukey's<br>post hoc) | Cohen's *d* | Interpretation |
| --- | --- | --- | --- | --- | --- |
| Envt. 1 | 0.001252 | Cont. vs. Low | 0.011 | 0.491 | Small |
|  |  | Cont. vs. Inter | 1.67E-04 | 1.043 | Large |
|  |  | Cont. vs. High | 1.67E-04 | 1.245 | Large |
|  |  | Low vs. Inter | 0.023 | 0.702 | Medium |
|  |  | Low vs. High | 0.002 | 0.997 | Large |
|  |  | Inter vs. High | 0.814 | 0.348 | Small |
| Envt. 2 | 5.68E-06 | Cont. vs. Low | 0.054 | 0.684 | Medium |
|  |  | Cont. vs. Inter | 1.75E-04 | 1.321 | Large |
|  |  | Cont. vs. High | 1.67E-04 | 2.330 | Large |
|  |  | Low vs. Inter | 0.037 | 0.626 | Medium |
|  |  | Low vs. High | 2.98E-04 | 1.302 | Large |
|  |  | Inter vs. High | 0.255 | 0.486 | Small |
| Envt. 3 | 5.07E-05 | Cont. vs. Low | 0.462 | 0.390 | Small |
|  |  | Cont. vs. Inter | 0.330 | 0.473 | Small |
|  |  | Cont. vs. High | 1.76E-04 | 1.915 | Large |
|  |  | Low vs. Inter | 0.995 | 0.059 | Small |
|  |  | Low vs. High | 0.002 | 1.184 | Large |
|  |  | Inter vs. High | 0.003 | 1.177 | Large |

**Table S4.** Summary of the *P*-values for Yield of clonal migration, post-selection. *P*-value of the main effect of migration was reported after Holm-Šidák correction for familywise error. Migration treatments were compared with each other using Tukey’s post-hoc analysis. Pairwise effect sizes were computed as Cohen’s *d*.

|  | Holm<br>Sidak<br>Corrected<br>*P*-values | Pairwise<br>comparison | *P*-values<br>(Tukey's<br>post hoc) | Cohen's *d* | Interpretation |
| --- | --- | --- | --- | --- | --- |
| Envt. 1 | 0.000116 | Cont. vs. Low | 0.171 | 0.598 | Medium |
|  |  | Cont. vs. Inter | 0.321 | 1.113 | Large |
|  |  | Cont. vs. High | 1.68E-04 | 1.304 | Large |
|  |  | Low vs. Inter | 0.984 | 0.642 | Medium |
|  |  | Low vs. High | 0.002 | 0.894 | Medium |
|  |  | Inter vs. High | 0.001 | 0.227 | Small |
| Envt. 2 | 1.11E-08 | Cont. vs. Low | 0.532 | 0.350 | Small |
|  |  | Cont. vs. Inter | 1.68E-04 | 1.763 | Large |
|  |  | Cont. vs. High | 1.67E-04 | 2.851 | Large |
|  |  | Low vs. Inter | 2.85E-04 | 1.088 | Large |
|  |  | Low vs. High | 1.67E-04 | 1.668 | Large |
|  |  | Inter vs. High | 0.441 | 0.465 | Small |
| Envt. 3 | 1.5E-05 | Cont. vs. Low | 0.034 | 0.567 | Medium |
|  |  | Cont. vs. Inter | 2.13E-04 | 0.479 | Small |
|  |  | Cont. vs. High | 1.72E-04 | 2.122 | Large |
|  |  | Low vs. Inter | 0.159 | 0.088 | Small |
|  |  | Low vs. High | 0.044 | 1.137 | Large |
|  |  | Inter vs. High | 0.934 | 1.271 | Large |

**Table S5.** Summary of the *P*-values for the growth rate of populations receiving migrants with variation. *P*-values of the main effect of migration was reported after Holm-Šidák correction for family-wise error. Migration treatments were compared with each other using Tukey’s post-hoc analysis. Pairwise effect sizes were computed as Cohen’s *d*.

|  | Holm<br>Sidak<br>Corrected<br>*P*-values | Pairwise<br>comparison | *P*-values<br>(Tukey's<br>post hoc) | Cohen's *d* | Interpretation |
| --- | --- | --- | --- | --- | --- |
| Env. 1 | 0.02406 | Cont. vs. Low | 0.984 | 0.109 | Small |
|  |  | Cont. vs. Inter | 0.612 | 0.395 | Small |
|  |  | Cont. vs. High | 0.032 | 0.847 | Large |
|  |  | Low vs. Inter | 0.819 | 0.283 | Small |
|  |  | Low vs. High | 0.074 | 0.747 | Medium |
|  |  | Inter vs. High | 0.377 | 0.510 | Medium |
| Env. 2 | 0.1435 | NA | NA | NA | NA |
| Env. 3 | 0.1296 | NA | NA | NA | NA |

**Table S6.** Summary of the *P*-values for yield of migration with variation. *P*-value of the main effect of migration was reported after Holm-Šidák correction for familywise error. Migration treatments were compared with each other using Tukey’s post-hoc analysis. Pairwise effect sizes were computed as Cohen’s d.

|  | Holm<br>Sidak<br>Corrected<br>*P*-values | Pairwise<br>comparison | *P*-values<br>(Tukey's<br>post hoc) | Cohen's *d* | Interpretation |
| --- | --- | --- | --- | --- | --- |
| Envt. 1 | 1.9E-05 | Cont. vs. Low | 0.581 | 0.528 | Medium |
|  |  | Cont. vs. Inter | 0.703 | 0.397 | Small |
|  |  | Cont. vs. High | 0.010 | 1.070 | Large |
|  |  | Low vs. Inter | 0.997 | 0.077 | Small |
|  |  | Low vs. High | 0.200 | 0.681 | Medium |
|  |  | Inter vs. High | 0.136 | 0.690 | Medium |
| Envt. 2 | 0.007 | Cont. vs. Low | 1.000 | 0.046 | Small |
|  |  | Cont. vs. Inter | 0.808 | 0.284 | Small |
|  |  | Cont. vs. High | 0.009 | 0.934 | Large |
|  |  | Low vs. Inter | 0.755 | 0.298 | Small |
|  |  | Low vs. High | 0.007 | 0.922 | Large |
|  |  | Inter vs. High | 0.084 | 0.555 | Medium |
| Envt. 3 | 0.010 | Cont. vs. Low | 0.680 | 0.371 | Small |
|  |  | Cont. vs. Inter | 0.328 | 0.587 | Medium |
|  |  | Cont. vs. High | 0.003 | 1.057 | Large |
|  |  | Low vs. Inter | 0.933 | 0.164 | Small |
|  |  | Low vs. High | 0.061 | 0.627 | Medium |
|  |  | Inter vs. High | 0.210 | 0.491 | Small |

